# Mutations in the riboflavin biosynthesis pathway confer resistance to furazolidone and abolish the synergistic interaction between furazolidone and vancomycin in *Escherichia coli*

**DOI:** 10.1101/2024.07.17.603971

**Authors:** Hannah Wykes, Vuong Van Hung Le, Jasna Rakonjac

## Abstract

Antibiotic combinations are a promising strategy to counteract the global problem of increasing antibiotic resistance. We have previously demonstrated furazolidone-vancomycin synergy against Gram-negative pathogens. Here, we selected *Escherichia coli* progeny for growth on the furazolidone-vancomycin combination to which the parent was sensitive. We show that selected clones were associated with increased resistance to neither, only one of, or both furazolidone and vancomycin, but in all cases were associated with a decrease in furazolidone-vancomycin synergy. Among a variety of gene mutations identified in this screen, we investigated the mechanism behind the most frequently arising mutations, those in the riboflavin biosynthesis genes *ribB* and *ribE*, and found them to act predominantly through decreasing the activity of the NfsA and NfsB nitroreductases, which have FMN (flavin mononucleotide) or FAD (flavin adenine dinucleotide) as a prosthetic group. We further show that the *ribB*/*ribE* mutants isolated in our screen are riboflavin semi-auxotrophs. Riboflavin supplementation restored the normal growth of the *ribB*/*ribE* mutants but not the furazolidone sensitivity.

## Introduction

Ever-increasing antibiotic resistance is a current and future global health issue; with the most urgent need identified by the World Health Organisation to develop treatments for Gram-negative bacteria. Among multiple strategies being developed, synergistic antibiotic combinations are clinically important for several reasons. Firstly, they lower the minimal effective dosage of each constituting drug, reducing side- effects and toxicity while broadening available drug options by including drugs that would otherwise be toxic at the effective dose in a mono-therapy (1). Secondly, synergistic combinations may be sufficient to kill mutants resistant to individual agents, suppressing the emergence of resistant mutants during combinatorial therapy (2). Nonetheless, the latter would be less significant if mutations arise that confer cross-resistance to both antibacterials and/or abolish the synergistic interaction. It is therefore important, following the discovery of a synergistic pair, to evaluate the emergence and phenotypes of the resistant mutants. Notably, isolating, identifying, and characterising mutations that cause synergy loss may reveal the molecular mechanism behind the interaction (3, 4).

We have previously reported that the combination of furazolidone, a nitrofuran antibiotic, and vancomycin, displays antibacterial synergy in *E. coli* (5). This combination holds promise to repurpose vancomycin, a high-molecular-weight glycopeptide antibiotic that poorly translocates across the outer membrane and is prescribed for the treatment of Gram-positive infections, into a treatment option for Gram-negative infections. In this work, we isolated and characterized mutations conferring resistance to the synergistic furazolidone-vancomycin combination, showing that the most frequent resistance mechanism was through the biosynthesis pathway of riboflavin, the precursor to the cofactors required for nitroreductases, enzymes responsible for furazolidone (prodrug) activation.

## Results

### Selecting antibacterial resistance mutations to the synergistic furazolidone-vancomycin combination

To isolate mutants resistant to the furazolidone-vancomycin combination, stationary-phase overnight cultures of BW25113 parental strain (PS) were spread on selective agar plates containing a combination of 256 mg/L vancomycin and 2 mg/L furazolidone (Supplementary Figure 1). Overall, seventeen resistant mutants were isolated and sequenced (Table 1). Different mutation types were found, including nonsense (*nlpI*), missense (*rpoC*), frameshift (*ftsH*, *wecC*, *opgG*), in-frame deletion (*ribE*), IS*1*/IS*5* insertions and point mutations in the 5’ untranslated region of *ribB*. Notably, most of the isolated resistant mutants were shown to contain mutations in essential genes: *ribB* (×4), *ribE* (×5), *ftsH* (×3), and *rpoC* (×1).

**Table 1.**
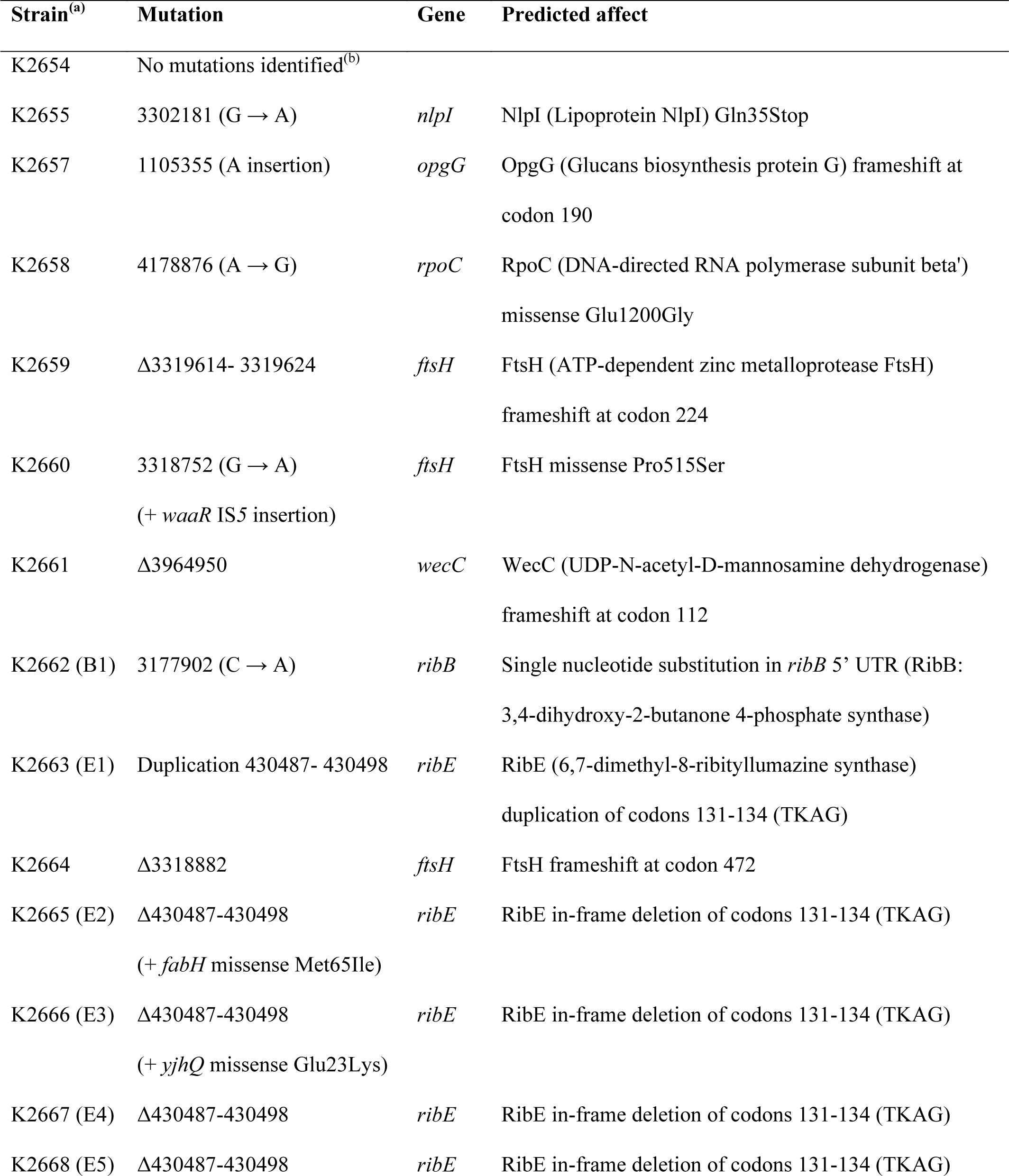
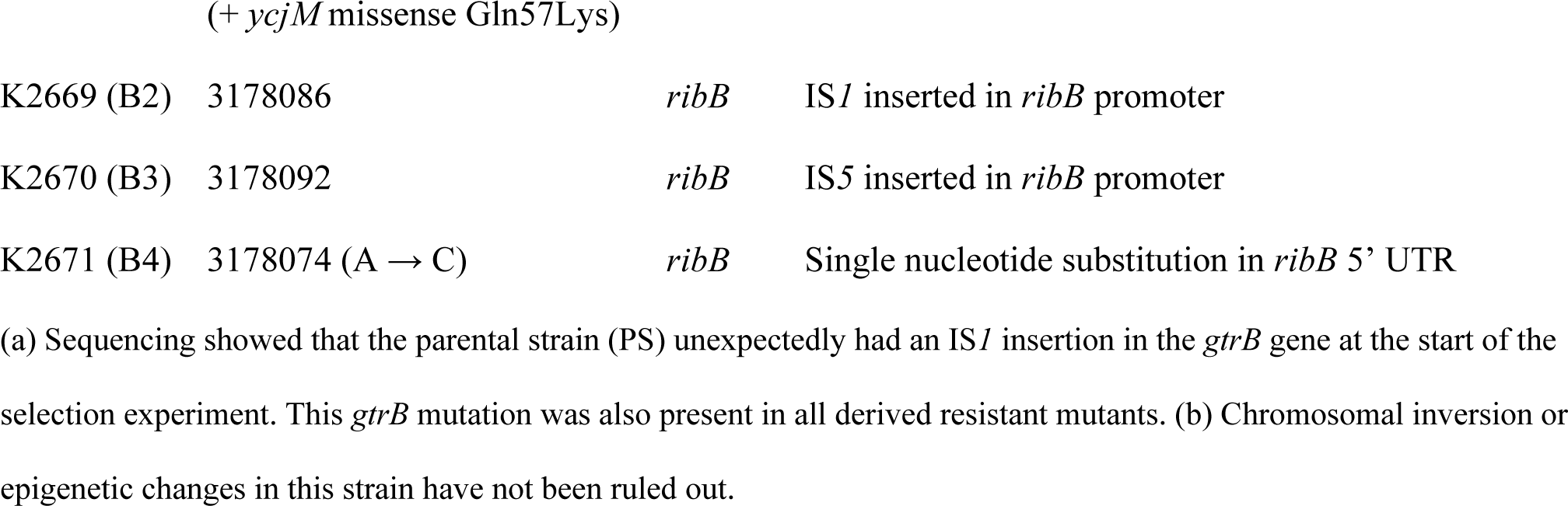
Mutations identified in isolated furazolidone-vancomycin resistant mutants in reference to the BW25113 genome (Accession number CP009273.1)

We also examined how individual antibiotic MICs and the furazolidone-vancomycin interaction changed in the isolated mutants, using antibiotic susceptibility broth microdilution and checkerboard assays, respectively. Strikingly, all isolated mutants demonstrated decreased synergy, whereas the changes in MICs for individual antibacterials fell into two main groups: I) Increased furazolidone resistance (with or without increased vancomycin resistance) and II) increased vancomycin resistance only (Figure 1). There was also one mutant which displayed decreased synergy with no individual MIC changes.

**Figure 1.**
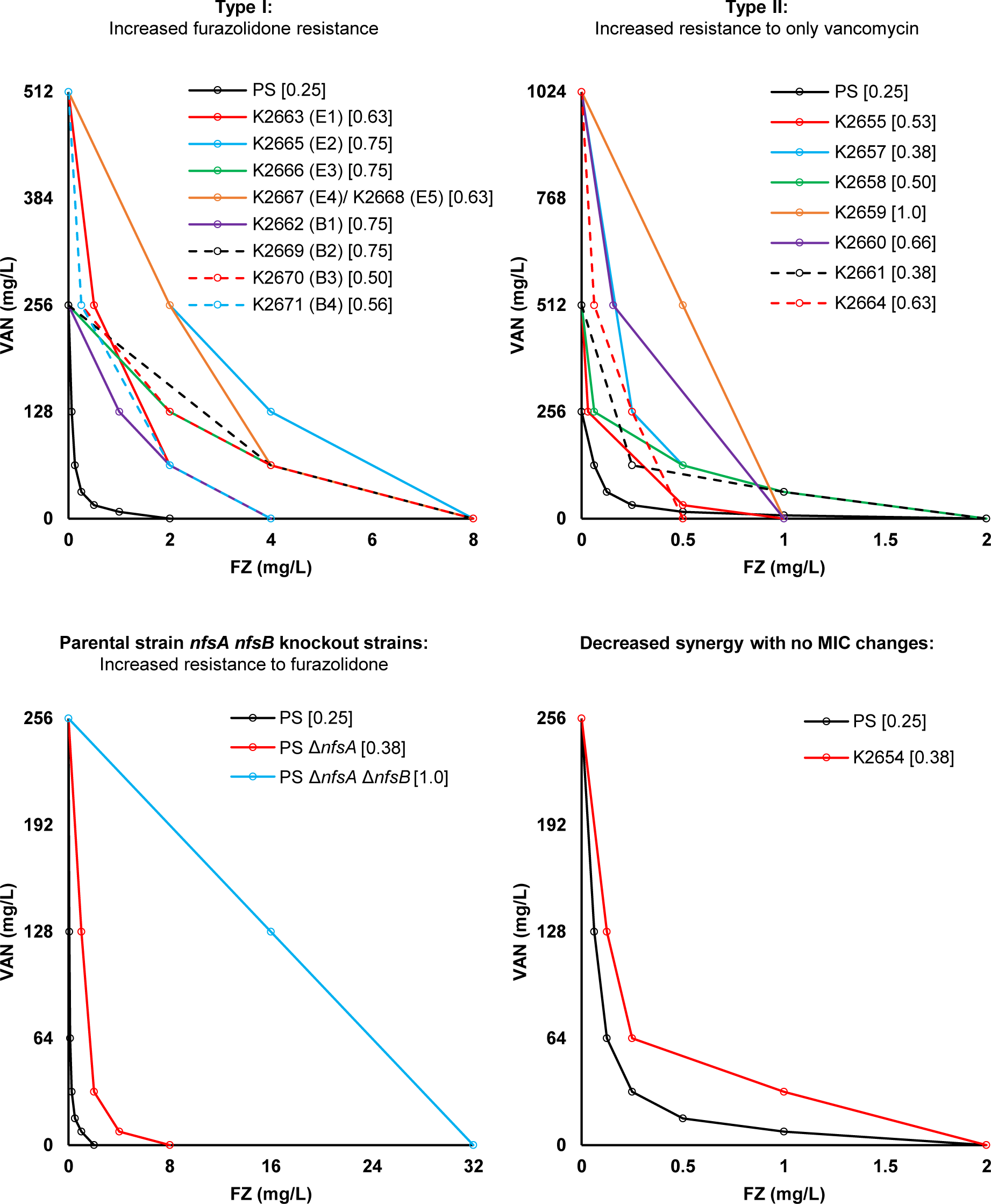
The vancomycin-furazolidone interaction in the isolated mutants. Checkerboard assays in liquid cultures were conducted on isolated resistant strains to construct the isobolograms for furazolidone- vancomycin interaction. All mutants had decreased synergy, reflected by their isobologram curve being less concave than the parental one. a) Type I, increased furazolidone resistance (includes resistance to both furazolidone and vancomycin). b) Type II, increased vancomycin resistance only. c) The Δ*nfsA* and Δ*nfsA* Δ*nfsB* mutations in the parental strain, which are known to confer nitrofuran resistance [23], but were not found in our double drug resistance selection experiment, are shown. d) The isobologram for K2654, which showed no MIC changes. Each point on the isobologram curve indicates the minimum concentration of each reagent in combination required to inhibit bacterial growth. The experiment was performed using three replicates, showing similar results. The FICI values for each strain are shown in square brackets. The *ribB* and *ribE* mutants are denoted by (B) and (E), respectively. PS, parental strain.

### Mutations in the riboflavin biosynthesis pathway are associated with furazolidone resistance

Of the seventeen isolated mutants, nine contained *ribB* or *ribE* mutations encoding enzymes in the riboflavin biosynthesis pathway (Supplementary Figure 2). This pathway is responsible for biosynthesis of FMN and FAD, cofactors for the two major nitrofuran-activating nitroreductases, NfsA and NfsB, and minor nitroreductase AhpF (6–8). These mutants demonstrated up to a four-fold increase in MIC_FZ_ (Figure 1). The *ribB* mutants all had mutations upstream of the coding region: B2 and B3 had IS*5* or IS*1* insertions in the promoter, and a 4-fold MIC_FZ_ increase, while B1 and B4 had single nucleotide substitutions in the 5’ untranslated region (5’ UTR) of the *ribB* mRNA (Figure 2a), and a two-fold MIC_FZ_ increase. The 5’-UTR of the *ribB* gene is a highly structured regulatory riboswitch that, upon binding flavin mononucleotide (FMN), represses *ribB* expression at both the transcriptional and translational levels (Figure 2b) (9).

**Figure 2.**
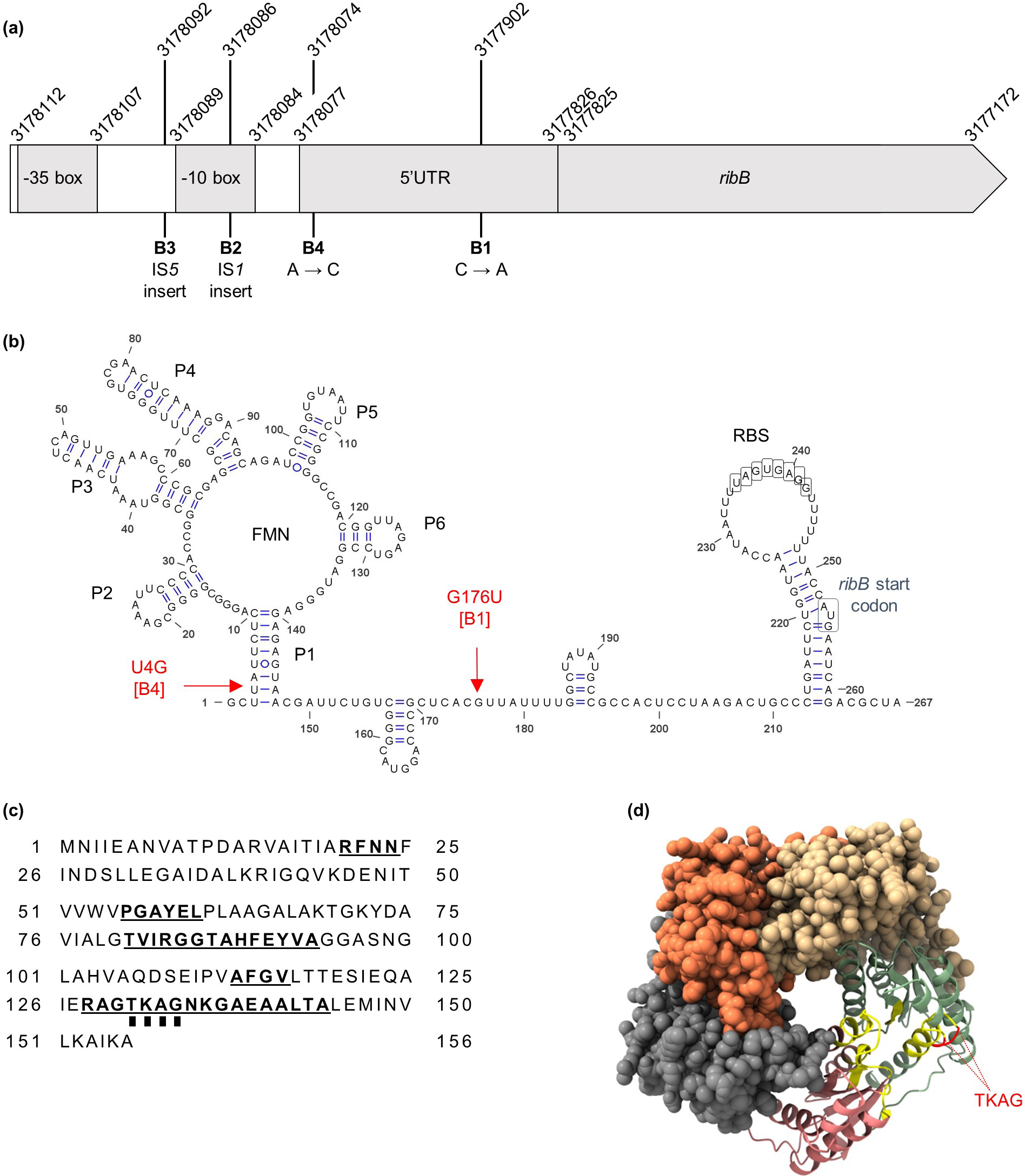
**Annotation of the *ribB*/*ribE* mutations**. (a) The position of the *ribB* mutations in strains B1, B2, B3 and B4. Mutants B1 and B4 had single nucleotide mutations in the mRNA 5’-untranslated region (5’-UTR), which forms a regulatory riboswitch. Mutants B2 and B3 had IS*1* and IS*5* insertions, respectively, within the promotor region. The genome coordinates are used in accordance with the BW25113 reference genome (GenBank accession number CP009273.1). Diagram not to scale. (b) Modelled secondary structure of the *ribB* riboswitch and the annotated mutations. (c) The amino acid sequence of the RibE protein. Residues making up the active site are emboldened and underlined [27]. The TKAG residues duplicated in the strain E1, and deleted in the strains E2, E3, E4, E5, are marked by squares underneath the residue letters. (d) The ColabFold-predicted model of a RibE pentamer. Each RibE biological complex is icosahedron composed of 60 monomeric units (= 12 pentamers). The active site residues are coloured yellow, and the mutated TKAG stretch is coloured red. RBS, ribosome binding site.

Regarding the *ribE* mutants, the same four amino acids (TKAG) were either deleted (mutants E2, E3, E4, E5) or duplicated (mutant E1) (Figure 2c, Table 1), causing a 4-fold or 2-fold increase in MIC_FZ_, respectively. We modelled a pentamer of the RibE icosahedron (10), showing that the TKAG residues are located at the interface of two adjacent monomeric subunits, in the active site of the complex (Figure 2d). Duplication or deletion of the TKAG residues is therefore expected to negatively affect the RibE enzyme activity.

### Growth rates and furazolidone dose-response curves of the *ribB*/*ribE* mutants

We next examined the *ribB*/*ribE* mutants’ growth in liquid broth. Most had noticeably slower growth than PS (Figure 3a & b). This was particularly severe in the *ribE* TKAG deletion mutants (E2, E3, E4, E5), which reached stationary phase earlier and at a much lower OD_600_ (∼0.2 vs ∼0.6) than E1, the TKAG duplication mutant.

**Figure 3.**
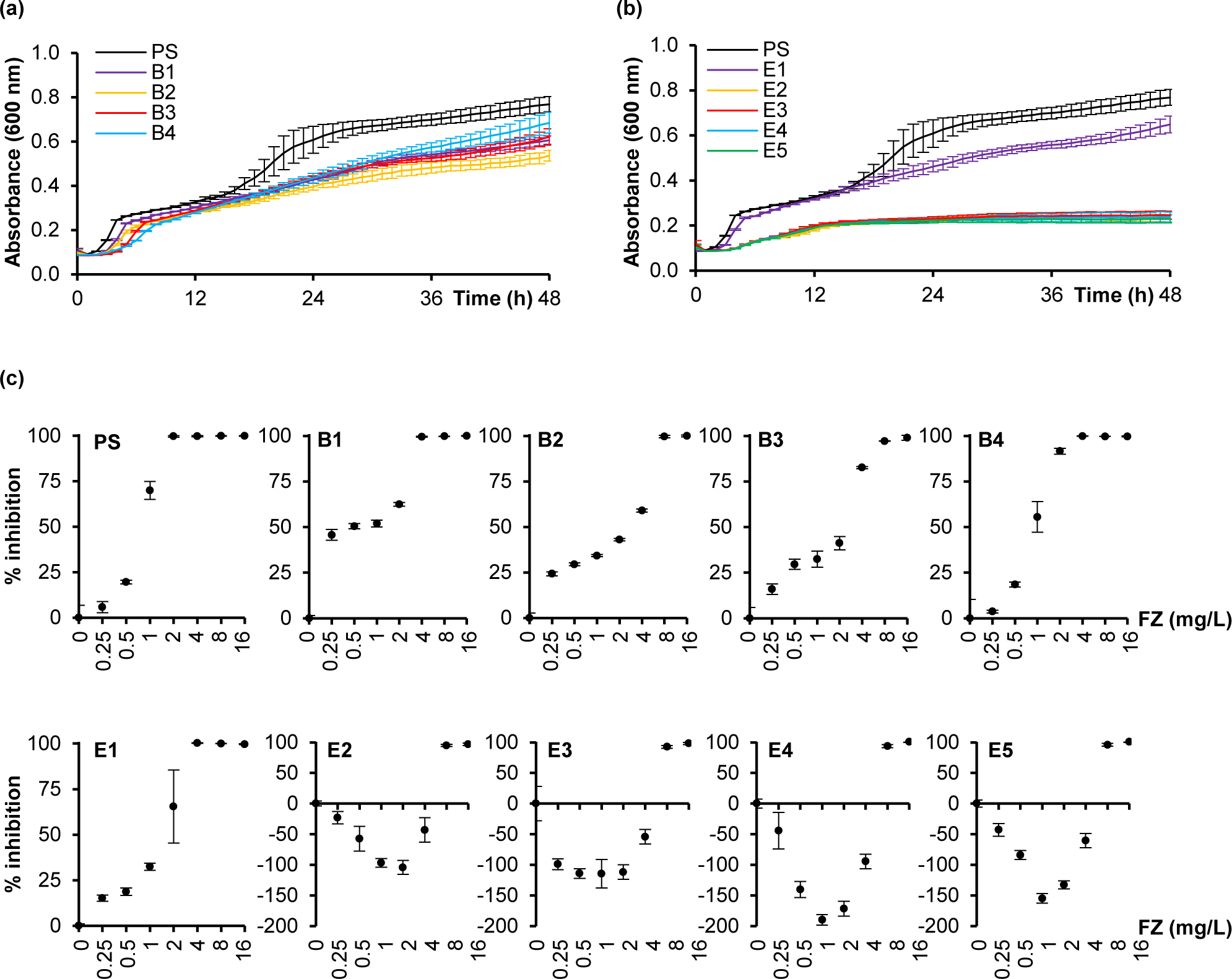
Growth and furazolidone dose-response inhibition profiles of the *ribB*/*ribE* mutants. Growth curves for the (a) *ribB* and (b) *ribE* mutants and parental strain (PS) were determined by measuring the OD_600_ every hour for 48 h. (c) Furazolidone (FZ) dose-response growth inhibition curves for the *ribB* and *ribE* mutants were determined by a broth microdilution assay at the 18 h timepoint. Growth inhibition was expressed as a percentage value of the antibiotic-containing culture O.D. relative to the cultures grown without antibiotic. Data shown is the mean ± standard deviation of three replicates.

In addition, furazolidone dose-response growth inhibition curves were performed to monitor the inhibitory effect of furazolidone concentration on growth (Figure 3c). The parental strain, all *ribB,* and the *ribE* TKAG duplication mutant E1, produced a typical sigmoidal dose-response inhibition curve. In contrast, a parabolic curve was observed for RibE TKAG deletion mutants E2, E3, E4 and E5, reflecting substantially improved growth at low furazolidone concentrations, peaking at 0.125 x MIC, with a 2- to 4-fold increased stationary phase OD_600_ relative to the no-furazolidone control.

### Complementation *in trans* reverses the furazolidone resistance and growth defect of the *ribB*/*ribE* **mutants**

We next asked if expressing the corresponding wild-type RibB/RibE proteins from ASKA collection plasmids (11) in the *ribB*/*ribE* mutants could lower the furazolidone MIC and restore the growth rate relative to the parental strain. Upon induction with 0.1 mM IPTG, the MIC_FZ_ was reduced to 1 mg/mL for the *ribB* mutants and 2 mg/mL for the *ribE* mutants, which is lower than, or equal to, the parental strain MIC_FZ_, respectively (Figure 4a). Notably, the MIC_FZ_ was decreased by 2-fold if the *ribB* or *ribE* gene was episomally expressed in PS (Figure 4a).

**Figure 4.**
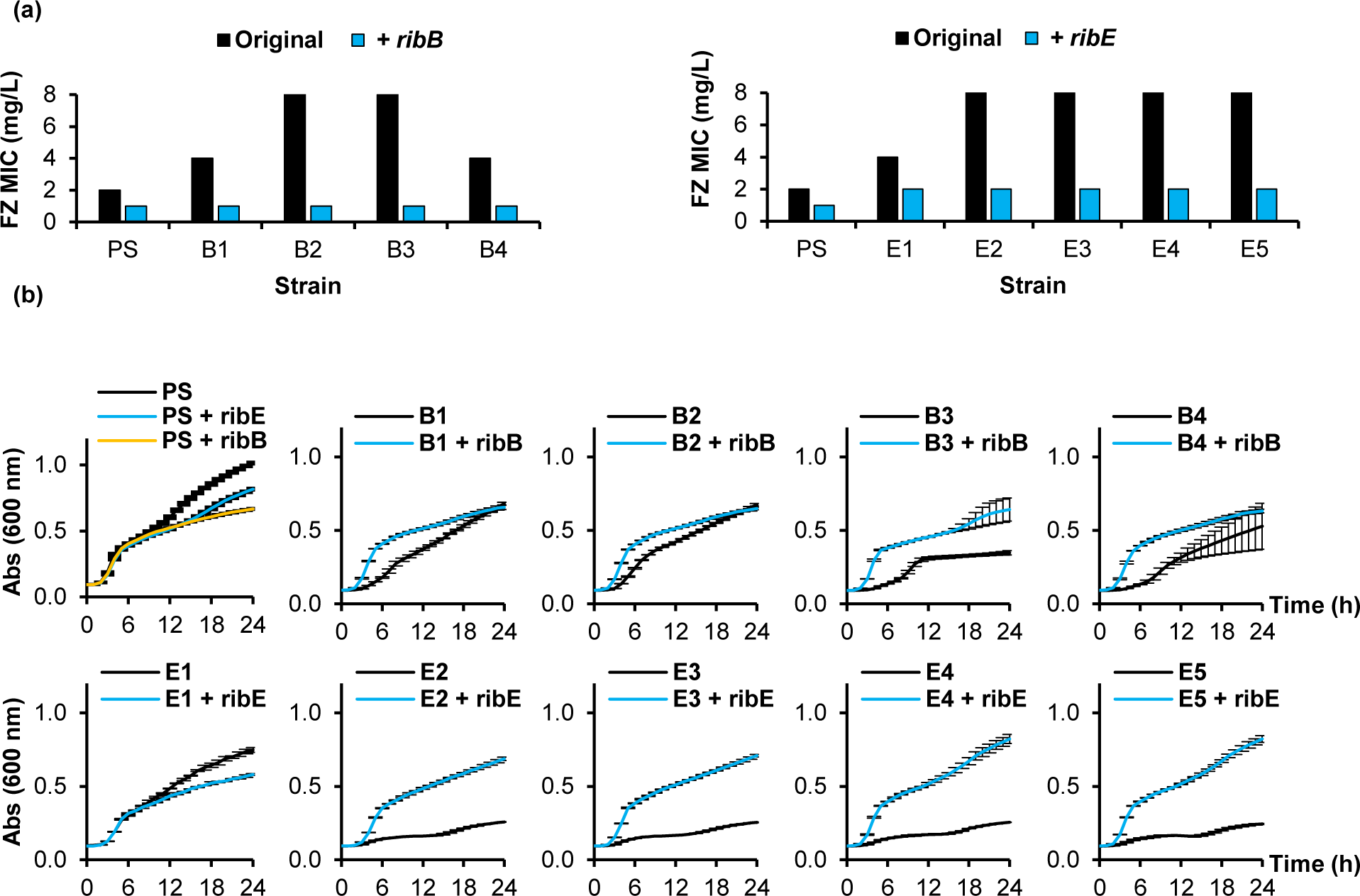
Furazolidone MICs and growth curves for the complemented *ribB*/*ribE* mutants. (a) The change in furazolidone (FZ) MIC upon complementation with either the functional *ribB* (PS, B1, B2, B3, B4) or *ribE* (PS, E1, E2, E3, E4, E5) gene. (b) Growth curves of the original and complemented strain at 37 °C. Expression was induced with 0.1 mM IPTG. Absorbance (Abs) at 600 nm was measured every hour for 24 h. Data shown is the mean ± standard deviation of three replicates.

Complementation with either *ribB* or *ribE* also improved the growth of all strains except the parental strain and E1, which had a much less severe growth impairment as compared to the other mutants (Figure 4b). Overall, complementation experiments confirm the causal role of the *ribB*/*ribE* mutations, rather than any secondary mutations identified (Table 1), for the furazolidone resistance and slow growth.

### The *ribB*/*ribE* mutations cause furazolidone resistance through decreasing the cellular furazolidone-activating nitroreductase activity

RibB and RibE are two essential enzymes in the biosynthesis pathway of riboflavin, the precursor for the cofactors (FMN, FAD) (9, 12) of the nitrofuran-activating nitroreductases (NfsA, NfsB, AhpF). To determine whether the *ribB*/*ribE* gene mutations affect the downstream nitroreductase activity, enzymatic assays were conducted on the cell extracts of PS and some representative isolated mutants; E1 (RibE TKAG duplication), E4 (RibE TKAG deletion, no secondary mutations), B2 and B3 (IS*1*/*5* insertion within the *ribB* promoter), as well as their corresponding complemented strains.

The cell lysate nitroreductase activities of the tested furazolidone-resistant mutants were lower than that of the parental strain (Figure 5a-c), indicating a lower furazolidone-activating rate. This enzymatic activity was increased to the parental strain equivalent when the mutants were complemented with the corresponding gene (*ribB* for B2/B3 or *ribE* for E1/E4). Noteworthily, this nitroreductase activity increase correlated with a MIC_FZ_ decrease in these complemented strains (Figure 5d). Taken together, the *ribB* and *ribE* mutations decreased cellular nitroreductase activity, which subsequently increased furazolidone resistance.

**Figure 5.**
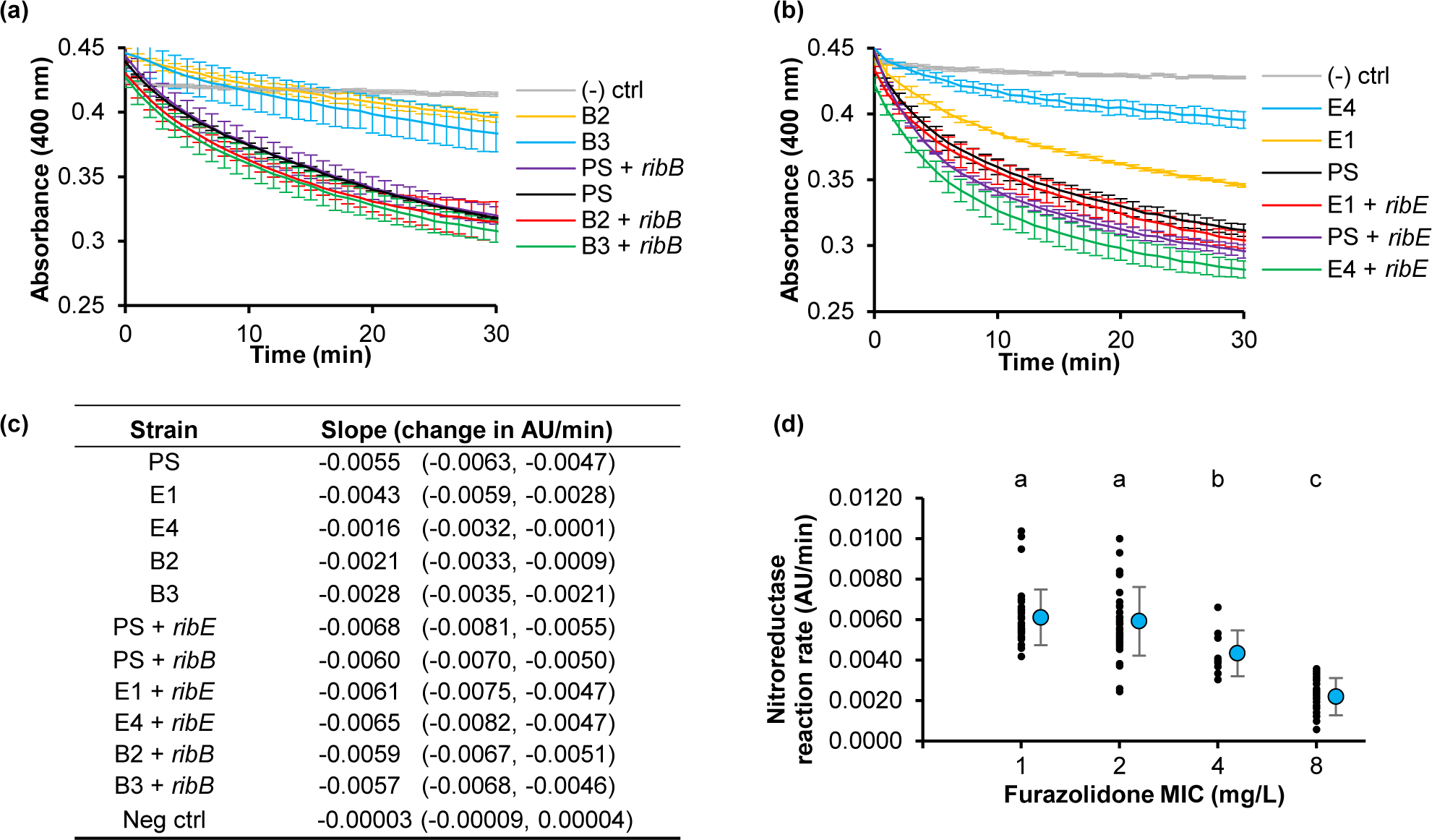
Nitroreductase assays for the *ribB*/*ribE* mutants and their complemented strains. a) Representative graphs showing the reaction progress curve for the nitroreductase assays for the *ribB* mutants and their corresponding complemented strains and (b) for the *ribE* mutants and their corresponding complemented strains. Each data point along the curve is the mean of three replicates ± standard deviation. Each reaction contained furazolidone, NADPH and the cellular lysate of the corresponding strain. The absorbance at 400 nm, indicating furazolidone concentration, was measured every minute for 12 h. (-) ctrl: negative control using the buffer in place of the cellular lysate. (c) The initial reaction velocity was calculated from three reaction replicates over the first ten minutes. The slope and 95% confidence interval are shown. (d) The correlation between the initial reaction velocity of the nitroreductase assays and the MIC_FZ_ of the furazolidone resistant mutants and the corresponding complemented strains. The mean and standard deviation for each MIC value is shown alongside each set of data points. Statistical difference between MIC groups was tested by One-Way ANOVA, followed by a Post-hoc Tukey-Kramer test. Different lowercase letters indicate a significant difference between any two MIC groups (p < 0.05); AU, arbitrary units.

### Effect of *nfsA*/*nfsB* knockout on furazolidone resistance in the *ribB*/*ribE* mutants

To determine whether the nitroreductase activity decrease was through the major nitroreductases NfsA and NfsB, Δ*nfsA* Δ*nfsB* double knockout strains were constructed in the *ribB*/*ribE* mutants and PS by sequential P1-mediated transduction and the MIC_FZ_ determined.

In the Δ*nfsA* Δ*nfsB* genetic background, the *ribB*/*ribE* strains were more than 2-fold closer in furazolidone MIC to the parental strain than in the wild-type *nfsA nfsB* background (Figure 6), indicating that the loss of *nfsA* and *nfsB* made the effect of the *ribB*/*ribE* mutations on furazolidone resistance redundant to some extent. Nonetheless, E4, still had increased furazolidone resistance in the Δ*nfsA* Δ*nfsB* genetic background. These findings suggest that the furazolidone resistance mediated by the *ribB*/*ribE* mutations was caused, though not entirely, through decreased NfsA/NfsB nitroreductase activity and that other factors may be involved in the furazolidone resistance.

**Figure 6.**
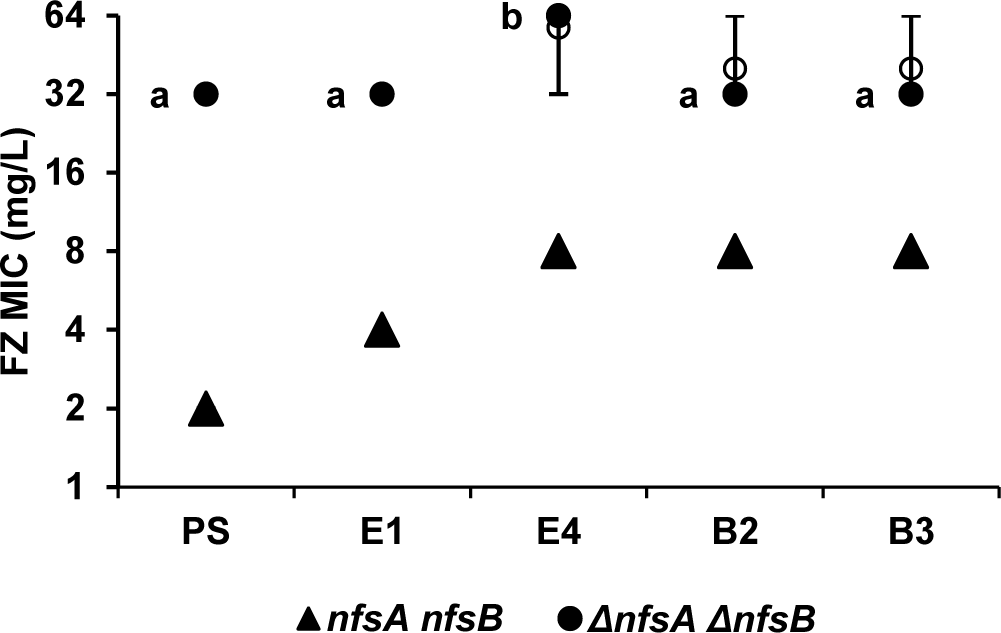
Effect of *nfsA*/*nfsB* knockout on the furazolidone MIC in the *ribB*/*ribE* mutants. Furazolidone MICs were obtained using standard broth microdilution assays. The strains tested were PS, and the E1, E4, B2 and B3 mutants containing wild-type *nfsA* and *nfsB* (solid triangles), and the Δ*nfsA* Δ*nfsB* knockout mutations (solid circles). At least four independent experiments were carried out for each strain. The range, median and mean are shown as bars, filled circles, and hollow circles, respectively. Statistical difference between MICs in the Δ*nfsA* Δ*nfsB* knockout mutation strains was tested by the Kruskal-Wallis test, followed by a Post-hoc Dunn’s test. Different lowercase letters indicate a significant difference between any two MIC groups (p < 0.05).

### Riboflavin supplementation enhances *ribB*/*ribE* mutant growth but does not affect the furazolidone sensitivity

Given that the *ribB*/*ribE* mutations decrease nitroreductase activity, probably *via* decreased efficiency in riboflavin biosynthesis (Supplementary Figure 2), the precursor of the nitroreductase cofactors (FMN/FAD), we hypothesised that exogenous addition of riboflavin could reverse the furazolidone resistance phenotype in the *ribB*/*ribE* mutants. The effect of 1 mM riboflavin supplementation was therefore investigated in the PS, E1, E4, B2, and B3 strains. We found that while growth was restored to that of PS, with all strains reaching an OD_600_ of around 0.7 at 24 hr (Figure 7b), all furazolidone MICs remained unchanged (Figure 7a). This rules out slow bacterial growth as a possible cause to the furazolidone resistance in the *ribB*/*ribE* mutants. Also, it shows that riboflavin supplementation is not viable as a strategy to re-sensitise the *ribB*/*ribE* mutants to furazolidone.

**Figure 7.**
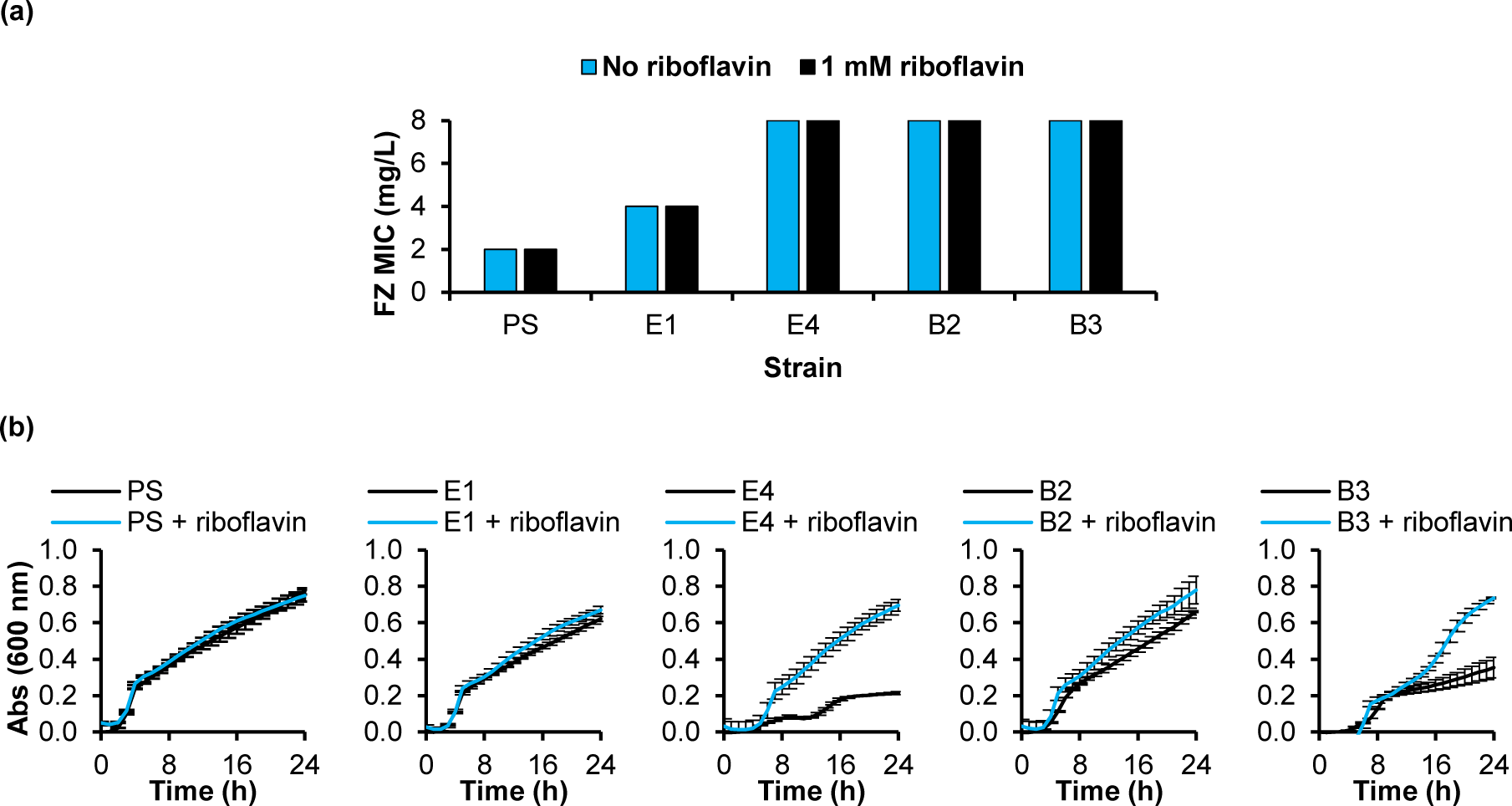
Effect of riboflavin supplementation on furazolidone sensitivity and growth. (a) Furazolidone (FZ) MICs and (b) growth curves of the furazolidone-resistant mutants and the parent strain upon riboflavin supplementation. Riboflavin was added at a concentration of 1 mM from the preparation of the overnight cultures. The absorbance at 600 nm was measured every hour for 24 h. Data shown is the mean ± standard deviation for three replicates.

### The TKAG deletion/duplication variants of RibE were found in *E. coli* multidrug resistant clinical isolates

We next asked if the *ribB*/*ribE* mutations in this study could be found in *E. coli* clinical isolates. Searching the RibE TKAG deletion and duplication variants against the NCBI genome database using Blastp (13) retrieved two and three clinical isolates for each mutant, respectively, some of which carry multiple antibiotic resistance genes, such as the strain BLSE9 from France and the strain E2010063_2015 from Australia (Table 2). By contrast, no clinical isolates were found to carry the *ribB* 5’-UTR nucleotide substitution or the promoter region IS*1/5* insertion mutations.

**Table 2:**
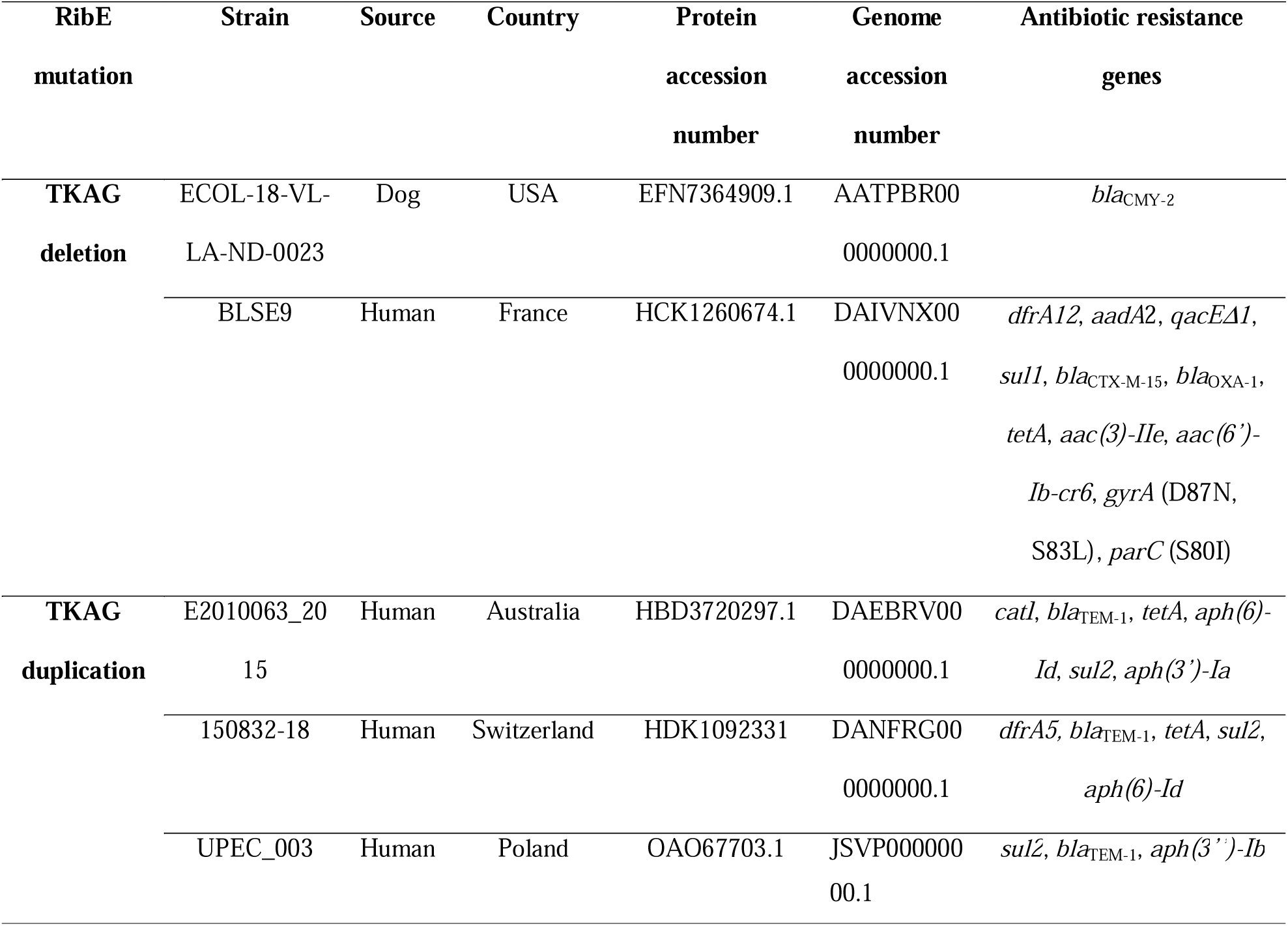
*E. coli* clinical isolates containing the RibE TKAG deletion or duplication mutations.

## Discussion

### Resistance to the furazolidone-vancomycin combination

We have previously shown furazolidone-vancomycin synergy against Gram-negative bacteria (5) and studied the bacterial response to this combination using transcriptomics (RNAseq) (14). In this work, we sought to further understand the synergy and potential resistance mechanisms to this combination by selecting and characterizing *E. coli* mutants isolated on furazolidone-vancomycin plates. This screen resulted in mutants with decreased synergy, divided into two groups: increased resistance to furazolidone through *ribB* and *ribE* mutations, or increased resistance to vancomycin (Figure 1).

Mutations in the *ftsH* gene were the most frequent amongst the increased vancomycin resistance group. Three different mutations of *ftsH* were isolated, all causing a loss of furazolidone-vancomycin synergy and having a collateral sensitivity phenotype (increased vancomycin resistance with increased furazolidone sensitivity) (Table 2, Figure 1). FtsH is an essential inner-membrane-anchored AAA^+^ protease that degrades specific proteinaceous targets for removal of misfolded proteins or regulated proteolysis in response to stresses (15). At least 23 FtsH substrates have been reported, including membrane-anchored and cytoplasmic targets, such as SecY, PspC, KdtA, LpxC, RpoH, SoxS, FolA and Cfa to name a few (15–17). It is very likely that the observed phenotypes are due to one or more of these FtsH substrates, whose identity remains to be determined. Future work is warranted to understand the role of the FtsH protein in the furazolidone-vancomycin synergy and collateral sensitivity to furazolidone.

### Mutations in the riboflavin biosynthesis pathway confer resistance to furazolidone

The largest proportion of mutants (9 of 17) had mutations in the essential *ribB* or *ribE* genes, which encode the RibB and RibE proteins in the riboflavin (vitamin B_2_) biosynthesis pathway (Supplementary Figure 2). Riboflavin is a precursor to FMN and FAD, cofactors required for the furazolidone-prodrug-activating nitroreductase enzymes NfsA, NfsB, and AhpF, in which the former two have a dominant role in drug activation. Using the nitroreductase assay, we established the correlation between the *ribB* and *ribE* mutations, the nitroreductase activity of the cellular lysate and the furazolidone resistance (Figure 5). The nitroreductase activity affected by the *ribB* and *ribE* mutations could predominantly be attributed to the two major nitroreductases, NfsA and NfsB. Deletion of *nfsA* and *nfsB* from the genomes of isolated *ribB* and *ribE* mutants and their analyses, however, still resulted in increased resistance in the E4 Δ*nfsA* Δ*nfsB* strain in comparison to the Δ*nfsA* Δ*nfsB* parent double mutant, pointing to additional furazolidone- activating enzymes, such as AhpF (8) or undiscovered ones, being involved (Figure 6).

It is worth mentioning the nature of the *ribB* and *ribE* mutations in this study. Since RibB and RibE are essential enzymes for *E. coli* survival, these mutations may decrease, but not totally abolish, the protein function. The *ribB* mutations were all upstream of the coding sequence, with mutants B2 and B3 having IS*1* and IS*5* insertions, respectively, in the promoter region and mutants B1 and B4 having point mutations in the 5’ UTR of the *ribB* mRNA (Figure 2). While it is reasonable to assume that disruptions to the promoter region would result in reduced transcription efficiency, how the mutations in the 5’-UTR lead to reduced RibB expression is less clear. The 5’-UTR of the *ribB* mRNA has been previously shown to form an FMN-binding riboswitch or aptamer (9) (Figure 2b). Binding of FMN to the aptamer prevents the formation of an anti-terminator/anti-sequester stem-loop, allowing the formation of a downstream terminator/ribosome binding site sequester stem-loop, inhibiting expression of *ribB* at both the transcriptional and translational level (9). Since the *ribB* mutations in the 5’-UTR found in the B1 and B4 isolates are associated with decreased RibB expression, supported by the increased resistance to furazolidone and restored sensitivity upon *ribB* complementation, these mutations must stabilise, not destabilise, the FMN-bound aptamer to further suppress the RibB translation.

RibE is an essential lumazine synthase in *E. coli* and is a hollow icosahedral complex composed of 60 subunits, assembled from 12 pentamers (18). All *ribE* mutations isolated here involved the same 12 nucleotides, encoding TKAG (codons 131-134). Mutant E1 had a TKAG duplication while mutants E2, E3, E4, and E5 had a TKAG deletion. These four residues are located in the interface between two adjacent monomers, involved in substrate binding (Figure 2c & d) (19), explaining why the enzymatic activity of the corresponding RibE mutant would be negatively impacted.

Notably, the same RibE TKAG deletion has been previously described, in an independent study, where it was selected by, and granted resistance to, nitrofurantoin, another nitrofuran antibiotic (20). This, and the fact that all the *ribE* mutants were independently isolated from separate plates in our screen, indicate that the *ribE* mutation to gain nitrofuran resistance is highly constrained and predictable.

In agreement with the essentiality of *ribB* and *ribE*, all mutants have shown slower growth than the parent, with the *ribE* TKAG deletion mutants being most affected. When riboflavin (metabolite downstream from the RibB and RibE catalysed reactions in the biosynthesis pathway) was supplemented in the medium, the growth defect was rectified. Most interestingly, however, riboflavin did not abolish furazolidone resistance, showing that slow bacterial growth has no role in the furazolidone resistance of the *ribB*/*ribE* mutants and ruling out the possibility of riboflavin supplementation to re-sensitise the *ribB*/*ribE* mutants to furazolidone. This observation likely reflects complex functional and regulatory roles of riboflavin. For example, riboflavin could be preferentially used by essential enzymes supporting bacterial growth, but not for functional restoration of the NfsA and NfsB enzymes. Another curiosity observed in this work is the growth-stimulatory effect of furazolidone at sublethal concentrations on the slow-growing *ribE* TKAG deletion mutants. This observation is in favour of direct activity of furazolidone as an electron donor or acceptor in essential biological processes that are normally dependent on FMN/FAD.

### Co-presence of furazolidone-resistant *ribE* mutations and other AMR genes in *E. coli* clinical isolates

Since the report of the *ribE* 12-nt deletion mutation in laboratory-selected nitrofurantoin resistant *E. coli* by Vervoort and colleagues (20), epidemiological studies have included the *ribE* gene besides the common targets, including *nfsA*, *nfsB* and *oqxAB*, when surveying nitrofurantoin resistance in clinical and environmental isolates (21–23). However, this *ribE* 12-nt TKAG^131-134^ deletion mutation has yet to be found in previous literature. One exception is the KAGN^132-135^ deletion in the RibE protein of the isolate EC0430U from the UK that overlaps with the TKAG^131-134^ deletion and was associated with increased nitrofurantoin resistance (22). By contrast, when searching the TKAG^131-134^ RibE variant against the NCBI database, we found two clinical isolates from the USA and France, where the latter also contains several other antibiotic resistance determinants (Table 2). Similarly, we found three *E. coli* clinical isolates containing the TKAG^131-134^ duplication with the co-occurrence of other AMR factors. The detection of these *ribE* mutations in clinical isolates, despite these mutations having significant fitness cost on the host, is concerning. This study provides evidence for three possible causes: i) the fitness cost can be compensated by external nutrients, such as riboflavin supplementation that improves the growth of the *ribE* mutants without re-sensitising the cell to furazolidone (Figure 7), ii) the *ribE* mutant may be co-selected with other AMR factors upon exposure to other antibiotics (Table 2), iii) the *ribE* mutant ‘feeds’ on furazolidone at sub-inhibitory concentrations via an unknown mechanism (Figure 3c). An alternative scenario is that compensatory mutations occur to improve the cell fitness through bypassing the decreased riboflavin biosynthesis pathway. Future work looking into this aspect of the *ribE* mutants is important to help devise a strategy to counter-select the nitrofuran-resistant *ribE* mutants.

In conclusion, we have shown that mutations affecting the *ribB* and *ribE* genes in the riboflavin biosynthesis pathway can confer resistance to the furazolidone-vancomycin combination through decreasing nitroreductase activity. In addition, these mutations were the most frequent in our screen, and mutations in the *ribE* gene have been previously reported as well as found in clinical isolates despite these mutations showing a significant fitness cost to the host in the absence of riboflavin.

## Materials and Methods

### Growth conditions and antibiotics

*E. coli* strains were grown at 37°C with shaking at 200 rpm. Growth media were either 2×YT (BD Difco^TM^) or CAMH (BD BBL^TM^) liquid broth, or solid plates (1% agar) (Pure Science). Antibiotics (GoldBio) stocks were made in water (ampicillin, kanamycin, vancomycin), or dimethyl sulfoxide (chloramphenicol, furazolidone).

### Bacterial strain construction

Bacterial strains and plasmids used in this study are shown in Table 3. Δ*nfsA* Δ*nfsB* double knock-outs of isolated mutants were constructed by stepwise rounds of P1 bacteriophage transduction (24) using single- gene knock-out mutations from the Keio collection as donors (25) followed by excision of the kanamycin resistance marker using FLP recombination as previously described (26). *E. coli* strains transformed with pCA24N and derived plasmids from ASKA collection (11) were grown in media containing 30 mg/L chloramphenicol and expression was induced with 0.1 mM IPTG, unless otherwise specified. Riboflavin was supplemented in the media at a final concentration of 1 mM.

**Table 3.**
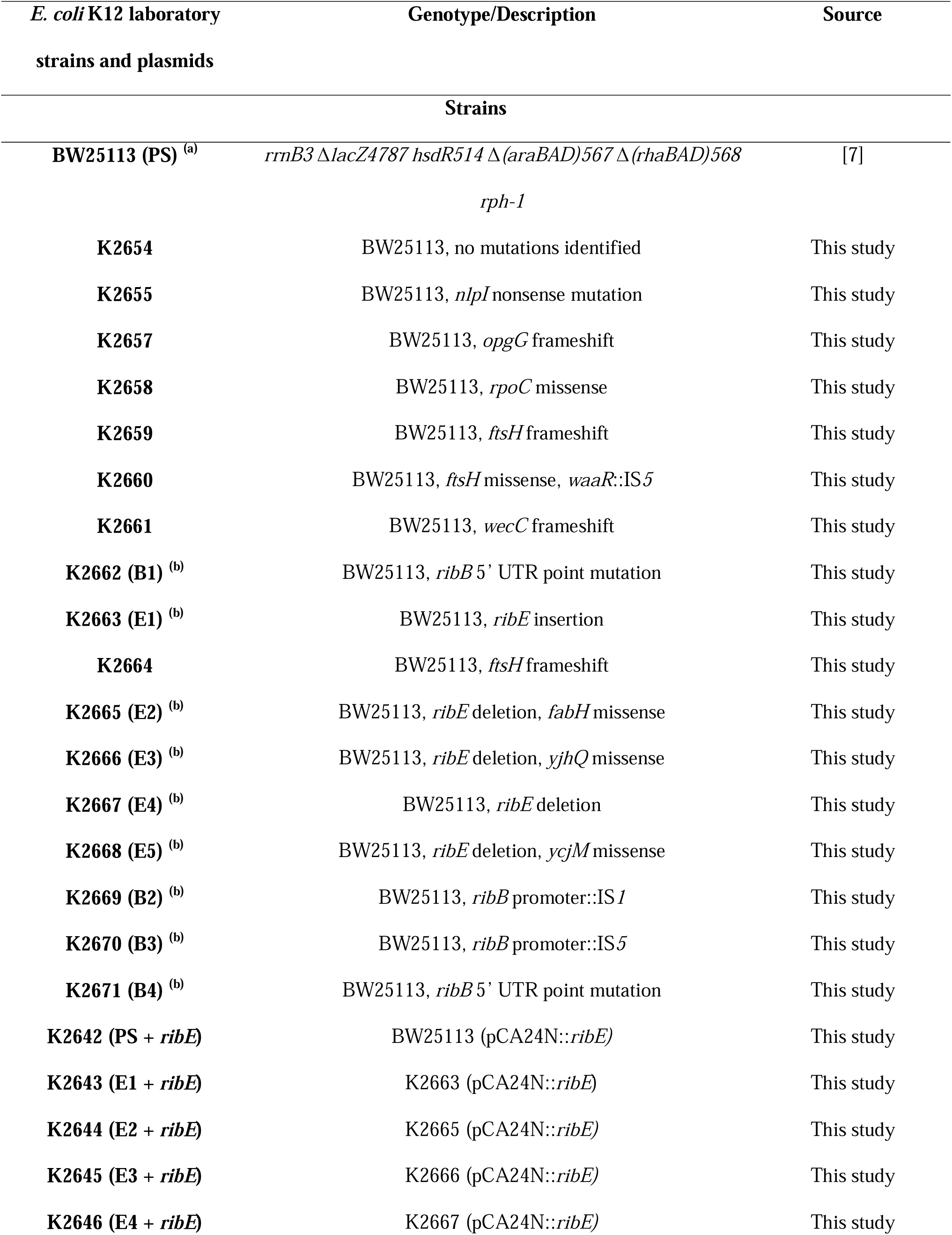
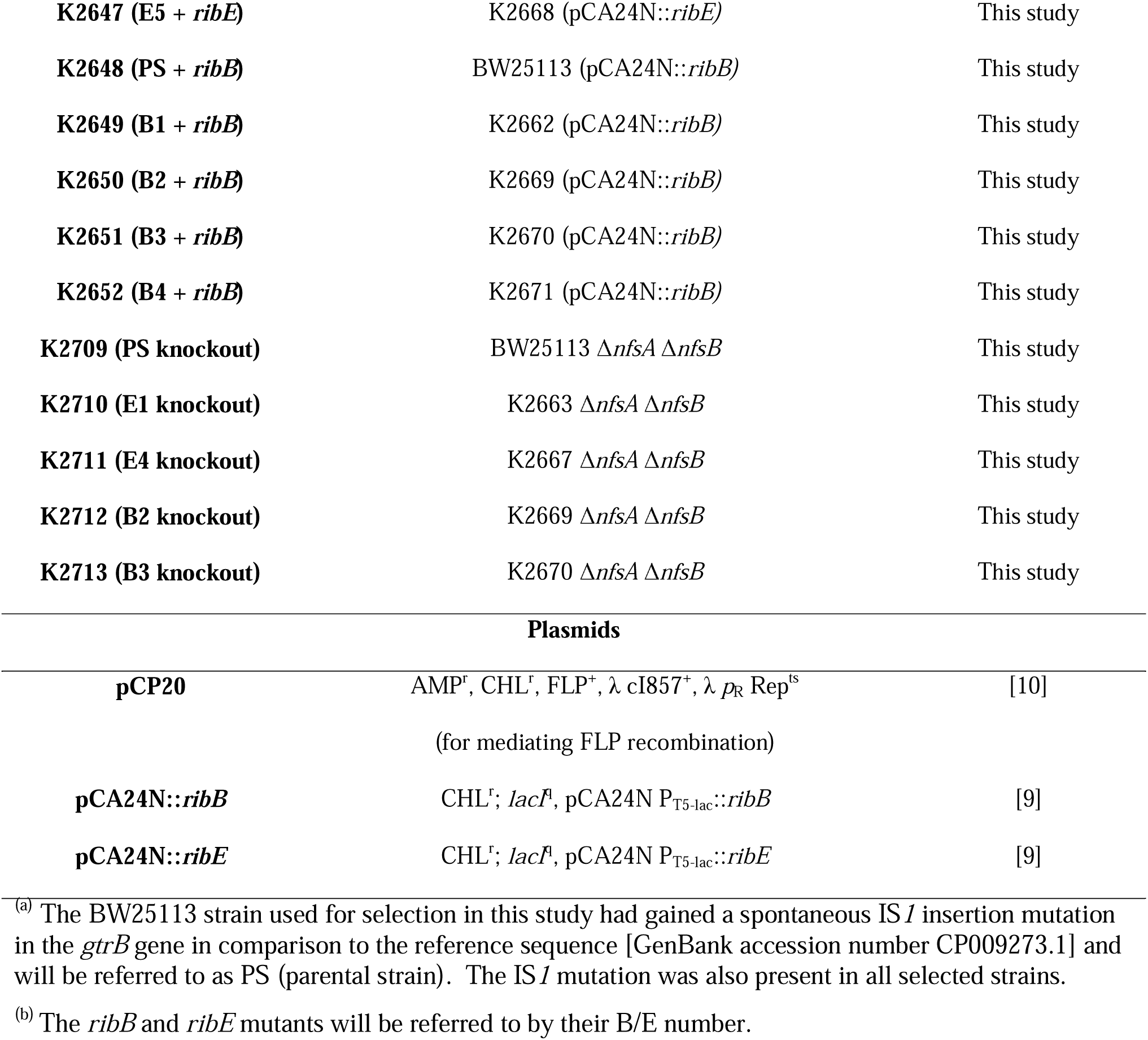
Bacterial strains and plasmids used in this study.

### Antimicrobial susceptibility assays and growth rate assays

Antibiotic MICs were determined according to CLSI guidelines (27) using broth microdilution and agar dilution methods. Growth rate assays were conducted as for the broth microdilution assays, with the optical density at 600 nm (OD_600_) measured every hour for either 24 or 48 h (Multiskan^TM^ GO Microplate Spectrophotometer).

### Growth inhibition checkerboard assays

Checkerboard assays were used to assess how the furazolidone-vancomycin interaction inhibits *E. coli* growth by standard microdilution method. Assays were conducted in CAMH broth in 384-well microplates. Two-fold serial dilutions of furazolidone and vancomycin were used. Each well contained 5×10^5^ cfu/mL, 1% DMSO, and antibiotics in a final volume of 50 μL. The microplates were incubated at 37 °C and the OD_600_ measured after 18 h (Multiskan^TM^ GO Microplate Spectrophotometer). Each treatment was performed in triplicate and the lowest drug concentration which caused a mean growth inhibition of at least 90 % in comparison to the no-antibiotic control was defined as MIC (28).

Fractional inhibitory concentration index (FICI) was calculated as follows:

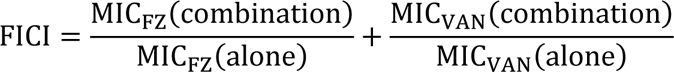

Where MIC_FZ_(combination) MIC_VAN_(combination) are the MICs for furazolidone and vancomycin when used in combination and MIC_FZ_(alone) MIC_VAN_(alone) are the MICs for furazolidone and vancomycin when used alone. The lowest FICI values were used to determine interactions: FICI ≤ 0.5 indicates synergy, FICI > 4 indicates antagonism, and 0.5 < FICI ≤ 4 indicates additivity (29).

### Isolating resistant mutants

Mutants of *E. coli* strain BW25113 were selected on CAMH agar containing a combination of vancomycin (256 mg/L) and furazolidone (2 mg/L). Twenty independent overnight cultures each inoculated from single colonies were separately spread on twenty selection plates. Briefly, 100 μL of each overnight culture was added to 2.5 mL of molten 0.5% CAMH agar (at ∼ 47 °C), vortexed, then poured onto the selective plate. Bacterial colonies were observed after 48 hours incubation, then sub-streaked onto non-selective agar plates. To minimise the chance of isolating colonies with identical mutations, only one colony was picked from each plate unless differences in colony morphology were observed.

### Comparative genome analysis

Genomic DNA was extracted using the DNeasy UltraClean Microbial Kit (Qiagen) according to the manufacturer’s instructions. The samples were submitted for whole genome sequencing to Massey Genome Service (Massey University, Palmerston North, New Zealand). Libraries were prepared using the Illumina DNA Prep kit and sequenced on the Illumina MiSeq™ 2×250-base paired-end v2 platform.

The raw reads were trimmed to an error probability cut-off of 0.001 (Phred score of 30), and reads less than 25 bases were removed using SolexaQA++ v3.1.7.1 (30). The trimmed reads were aligned to the reference genome (*E. coli* BW25113 accession number CP009273.1) (31) using bowtie2 v2.4.2 (32) in the --very-sensitive mode. SAMtools v1.14 (33) was used to convert the SAM sequence alignment files into BAM files, followed by variant calling using freebayes v1.3.1 (34), with the ploidy set to 1. The variants were annotated using SnpEff v4.4.20(1) (35).

Genomic structural variations were identified by extracting the unmapped reads using SAMtools v1.14 (33), which were then assembled into contigs using SPAdes v3.13.0 (36), using the careful mode. The generated contigs were mapped to the reference *E. coli* BW25113 genome using National Center for Biotechnology Information (NCBI) nucleotide BLAST+ 2.12.0 (13) to determine the location of any structural variations, if present.

### RNA and protein modelling

The homology-based secondary structural model of the FMN aptamer at 5’-untranslated region of the *ribB* mRNA (corresponding to the reverse strand at the coordinates 3177808-3178077 of the reference *E. coli* BW25113 genome) was extracted from (37) and visualized using Varne v3.9 (38). One of the 12 pentamers that form the 60-subunit RibE icosahedral biological complex was modelled using ColabFold v1.3.0 with default parameters (10) with the input being five copies of the RibE amino acid sequence (GenBank accession number AIN30914.1) separated by colons.

### Nitroreductase activity assays

Nitroreductase activity assays (8) were conducted on cell extracts of selected *ribB*/*ribE* mutants, PS, and the corresponding *ribB*/*ribE* complemented strains. Each strain was analysed in three independent assays.

Overnight cultures were diluted 1:100 into 25 mL of CAMH broth and grown to OD_600_ ∼0.5 at 37°C, centrifuged (10 min, 4000 x g), and the pellets stored at -20°C until use. The pellets were washed with 10 mL of pre-chilled 50 mM Tris-HCl (pH 7.4), centrifuged (10 min, 4000 x g, 4°C), and resuspended in 3.5 mL of pre-chilled 50 mM Tris-HCl (pH 7.4). The OD_600_ of each cell suspension was measured and adjusted with 50 mM Tris-HCl to a concentration of 1×10^9^ cfu/mL. Next, 3 mL of this cell suspension was sonicated (amplitude 15 for 4 min, 2 seconds on, 2 seconds off) using the microtip of a Virsonic 600 ultrasonic cell disruptor (Qsonica). The cell lysate was then centrifuged (14,000 x g, 30 min, RT) and the supernatant was collected for enzymatic analyses.

Nitroreductase activity assays were performed on a 96-well plate and each reaction was performed in triplicate. Each well contained 0.1 mM NADPH (Roche), 0.1 mM furazolidone, and 50 μL cell-extract in 50 mM Tris-HCl (pH 7.4) in a total volume of 200 μL. NADPH was added last to initiate the reaction.

Wells without cell-extract were used as negative controls. The assay was incubated at 25 °C, and absorbance at 400 nm was measured every minute for 12 h.

### Searching for the *ribB*/*ribE* mutations in clinical isolates

For the TKAG deletion/duplication mutations found in the *ribE* mutants, the corresponding RibE amino acid sequence (GenBank accession no. AIN30914.1 with TKAG deletion/duplication) was queried against the NCBI non-redundant protein sequence using the Blastp webserver (13). For the mutation in the 5’-untranslated region of *ribB*, the corresponding mutated nucleotide sequence ranging from 3177808- 3178077 of the reference genome BW25113 was queried against the GenBank nucleotide collection using megaBlast with default parameters (39). For the insertional mutation within the promoter of the *ribB* gene, an *in-silico* PCR was used. A pair of primers targeting the *ribB* promoter was designed using the primerBlast webserver (40), 5’-GGTTACCAGAATCAGGGCAGT-3’ and 5’- GTTGAGTGCCATTGTAGTGCG-3’, and then queried using the same tool with default parameters except setting *Escherichia coli* as the searching database to predict the amplicon size. The amplicon size of the wildtype was predicted to be 324 bp while the mutants containing IS*1*/IS*5* within the *ribB* promoter were expected to have a larger amplicon by 0.8-1.2 kb. Noteworthily, this *in silico* PCR would not detect the transpositional mutations for incomplete fragmented genome assemblies. If any *E. coli* isolate containing IS*1*/IS*5* transposition within the *ribB* promoter in the database was sequenced and assembled with short-read sequencing technique only, the genome assembly would be fragmented at the insertional site due to the presence of multiple copies of the IS*1*/IS*5* elements in a genome and therefore the *in silico* PCR would fail to generate a correct amplicon.

The bacterial genome assembly containing the queried mutation was retrieved and searched against the Comprehensive Antibiotic Resistance Database (CARD) with default parameters to identify the presence of other AMR genes (41).

### NCBI GenBank accession numbers

The raw sequencing reads are available from the NCBI Sequence Read Archive under BioProject accession PRJNA854676. (The parental strain BW25113 is called K2653 in the sequencing data). https://www.ncbi.nlm.nih.gov/sra/?term=PRJNA854676

## Acknowledgements

This work was supported by a Massey University-MBIE PSAF II grant MU001985 and a generous donation by Anne and Bryce Carmine as well as the Massey University School of Natural Sciences. H.W. was supported by the Graduate Women Manawatū Charitable Trust and the William Georgetti Scholarship.

## Transparency declaration

None to declare.

## Notes

### Competing Interest Statement

The authors have declared no competing interest.

### Summary of Updates

To add the Supplementary Materials file

## References

1. Sun W, Sanderson PE, Zheng W. 2016. Drug combination therapy increases successful drug repositioning. Drug Discovery Today 21:1189–1195.

2. Urban C, Mariano N, Rahal JJ. 2010. *In vitro* double and triple bactericidal activities of doripenem, polymyxin B, and rifampin against multidrug-resistant *Acinetobacter baumannii*, *Pseudomonas aeruginosa*, *Klebsiella pneumoniae*, and *Escherichia coli*. Antimicrobial Agents and Chemotherapy 54:2732–2734.

3. Chevereau G, Bollenbach T. 2015. Systematic discovery of drug interaction mechanisms. Molecular Systems Biology 11:807.

4. Le VVH, Olivera C, Spagnuolo J, Davies IG, Rakonjac J. 2020. *In vitro* synergy between sodium deoxycholate and furazolidone against enterobacteria. BMC Microbiology 20:5.

5. Olivera C, Le VVH, Davenport C, Rakonjac J. 2021. *In vitro* synergy of 5-nitrofurans, vancomycin and sodium deoxycholate against Gram-negative pathogens. Journal of Medical Microbiology 70.

6. Zenno S, Koike H, Kumar AN, Jayaraman R, Tanokura M, Saigo K. 1996. Biochemical characterization of NfsA, the *Escherichia coli* major nitroreductase exhibiting a high amino acid sequence homology to Frp, a *Vibrio harveyi* flavin oxidoreductase. Journal of bacteriology 178:4508–4514.

7. Zenno S, Koike H, Tanokura M, Saigo K. 1996. Gene cloning, purification, and characterization of NfsB, a minor oxygen-insensitive nitroreductase from *Escherichia coli*, similar in biochemical properties to FRase I, the major flavin reductase in *Vibrio fischeri*. The Journal of Biochemistry 120:736–744.

8. Le VVH, Davies IG, Moon CD, Wheeler D, Biggs PJ, Rakonjac J. 2019. Novel 5-nitrofuran- activating reductase in *Escherichia coli*. Antimicrobial Agents and Chemotherapy 63:e00868–19.

9. Pedrolli D, Langer S, Hobl B, Schwarz J, Hashimoto M, Mack M. 2015. The *ribB* FMN riboswitch from *Escherichia coli* operates at the transcriptional and translational level and regulates riboflavin biosynthesis. The FEBS journal 282:3230–3242.

10. Mirdita M, Schütze K, Moriwaki Y, Heo L, Ovchinnikov S, Steinegger M. 2022. ColabFold: making protein folding accessible to all. Nature Methods 19:679–682.

11. Kitagawa M, Ara T, Arifuzzaman M, Ioka-Nakamichi T, Inamoto E, Toyonaga H, Mori H. 2005. Complete set of ORF clones of *Escherichia coli* ASKA library (A complete set of *E. coli* K-12 ORF archive): Unique resources for biological research. DNA research 12:291–299.

12. Averianova LA, Balabanova LA, Son OM, Podvolotskaya AB, Tekutyeva LA. 2020. Production of vitamin B2 (riboflavin) by microorganisms: An overview. Frontiers in Bioengineering and Biotechnology 8.

13. Altschul SF, Madden TL, Schäffer AA, Zhang J, Zhang Z, Miller W, Lipman DJ. 1997. Gapped BLAST and PSI-BLAST: a new generation of protein database search programs. Nucleic Acids Research 25:3389–3402.

14. Olivera C, Cox MP, Rowlands GJ, Rakonjac J. 2021. Correlated transcriptional responses provide insights into the synergy mechanisms of the furazolidone, vancomycin, and sodium deoxycholate triple combination in *Escherichia coli*. mSphere 6:e00627-21.

15. Bittner L-M, Arends J, Narberhaus F. 2017. When, how and why? Regulated proteolysis by the essential FtsH protease in *Escherichia coli*. Biological Chemistry 398:625–635.

16. Hari SB, Morehouse JP, Baker TA, Sauer RT. 2023. FtsH degrades kinetically stable dimers of cyclopropane fatty acid synthase via an internal degron. Molecular Microbiology 119:101–111.

17. Morehouse JP, Baker TA, Sauer RT. 2022. FtsH degrades dihydrofolate reductase by recognizing a partially folded species. Protein Science 31:e4410.

18. Mörtl S, Fischer M, Richter G, Tack J, Weinkauf S, Bacher A. 1996. Biosynthesis of riboflavin. Lumazine synthase of Escherichia coli. J Biol Chem 271:33201–7.

19. Ritsert K, Huber R, Turk D, Ladenstein R, Schmidt-Bäse K, Bacher A. 1995. Studies on the Lumazine Synthase/Riboflavin Synthase Complex ofBacillus subtilis: Crystal Structure Analysis of Reconstituted, Icosahedral β-subunit Capsids with Bound Substrate Analogue Inhibitor at 2.4 Å Resolution. Journal of Molecular Biology 253:151-167.

20. Vervoort J, Xavier BB, Stewardson A, Coenen S, Godycki-Cwirko M, Adriaenssens N, Kowalczyk A, Lammens C, Harbarth S, Goossens H. 2014. An *in vitro* deletion in *ribE* encoding lumazine synthase contributes to nitrofurantoin resistance in *Escherichia coli*. Antimicrobial agents and chemotherapy 58:7225–7233.

21. Ho P-L, Ng K-Y, Lo W-U, Law PY, Lai EL-Y, Wang Y, Chow K-H. 2016. Plasmid-mediated OqxAB is an important mechanism for nitrofurantoin resistance in *Escherichia coli*. Antimicrobial Agents and Chemotherapy 60:537–543.

22. Wan Y, Mills E, Leung RCY, Vieira A, Zhi X, Croucher NJ, Woodford N, Jauneikaite E, Ellington MJ, Sriskandan S. 2021. Alterations in chromosomal genes *nfsA*, *nfsB*, and *ribE* are associated with nitrofurantoin resistance in *Escherichia coli* from the United Kingdom. Microbial Genomics 7.

23. Khamari B, Adak S, Chanakya PP, Lama M, Peketi ASK, Gurung SA, Chettri S, Kumar P, Bulagonda EP. 2022. Prediction of nitrofurantoin resistance among Enterobacteriaceae and mutational landscape of *in vitro* selected resistant *Escherichia coli*. Research in Microbiology 173:103889.

24. Thomason L, Constantino N. 2007. Court DL *E. coli* manipulation by P1 transduction. Current Protocols in Molecular Biology.

25. Baba T, Ara T, Hasegawa M, Takai Y, Okumura Y, Baba M, Datsenko KA, Tomita M, Wanner BL, Mori H. 2006. Construction of *Escherichia coli* K 12 in frame, single gene knockout mutants: the Keio collection. Molecular systems biology 2:2006.0008.

26. Datsenko KA, Wanner BL. 2000. One-step inactivation of chromosomal genes in *Escherichia coli* K-12 using PCR products. Proceedings of the National Academy of Sciences 97:6640–6645.

27. Cockerill FR, Wikler MA, Alder J, Dudley MN, Eliopoulos GM, Ferraro MJ, Hardy DJ, Hecht DW, Hindler JA, Patel JB, Powell M, Swenson JM, Richard B. Thomson J, Traczewski MM, Turnidge JD, Weinstein MP, Zimmer BL. 2012. Methods for dilution antimicrobial susceptibility tests for bacteria that grow aerobically; Approved Standard—Ninth Edition. Clinical and Laboratory Standards Institute,

28. Campbell J. 2010. High-Throughput Assessment of Bacterial Growth Inhibition by Optical Density Measurements. Current Protocols in Chemical Biology 2:195–208.

29. Odds FC. 2003. Synergy, antagonism, and what the chequerboard puts between them. Journal of Antimicrobial Chemotherapy 52:1–1.

30. Cox MP, Peterson DA, Biggs PJ. 2010. SolexaQA: At-a-glance quality assessment of Illumina second-generation sequencing data. BMC bioinformatics 11:1–6.

31. Grenier F, Matteau D, Baby V, Rodrigue S. 2014. Complete genome sequence of *Escherichia coli* BW25113. Genome Announcements 2.

32. Langmead B, Salzberg SL. 2012. Fast gapped-read alignment with Bowtie 2. Nature methods 9:357–359.

33. Li H, Handsaker B, Wysoker A, Fennell T, Ruan J, Homer N, Marth G, Abecasis G, Durbin R. 2009. The sequence alignment/map format and SAMtools. Bioinformatics 25:2078–2079.

34. Garrison E, Marth G. 2012. Haplotype-based variant detection from short-read sequencing. arXiv preprint arXiv:12073907.

35. Cingolani P, Platts A, Wang LL, Coon M, Nguyen T, Wang L, Land SJ, Lu X, Ruden DM. 2012. A program for annotating and predicting the effects of single nucleotide polymorphisms, SnpEff: SNPs in the genome of *Drosophila melanogaster* strain w1118; iso-2; iso-3. Fly 6:80-92.

36. Bankevich A, Nurk S, Antipov D, Gurevich AA, Dvorkin M, Kulikov AS, Lesin VM, Nikolenko SI, Pham S, Prjibelski AD. 2012. SPAdes: a new genome assembly algorithm and its applications to single-cell sequencing. Journal of computational biology 19:455–477.

37. Howe JA, Wang H, Fischmann TO, Balibar CJ, Xiao L, Galgoci AM, Malinverni JC, Mayhood T, Villafania A, Nahvi A, Murgolo N, Barbieri CM, Mann PA, Carr D, Xia E, Zuck P, Riley D, Painter RE, Walker SS, Sherborne B, de Jesus R, Pan W, Plotkin MA, Wu J, Rindgen D, Cummings J, Garlisi CG, Zhang R, Sheth PR, Gill CJ, Tang H, Roemer T. 2015. Selective small-molecule inhibition of an RNA structural element. Nature 526:672–677.

38. Darty K, Denise A, Ponty Y. 2009. VARNA: Interactive drawing and editing of the RNA secondary structure. Bioinformatics 25:1974-1975.

39. Morgulis A, Coulouris G, Raytselis Y, Madden TL, Agarwala R, Schäffer AA. 2008. Database indexing for production MegaBLAST searches. Bioinformatics 24:1757–64.

40. Ye J, Coulouris G, Zaretskaya I, Cutcutache I, Rozen S, Madden TL. 2012. Primer-BLAST: A tool to design target-specific primers for polymerase chain reaction. BMC Bioinformatics 13:134.

41. Alcock BP, Huynh W, Chalil R, Smith KW, Raphenya AR, Wlodarski MA, Edalatmand A, Petkau A, Syed SA, Tsang KK, Baker SJC, Dave M, McCarthy MC, Mukiri KM, Nasir JA, Golbon B, Imtiaz H, Jiang X, Kaur K, Kwong M, Liang ZC, Niu KC, Shan P, Yang JYJ, Gray KL, Hoad GR, Jia B, Bhando T, Carfrae LA, Farha MA, French S, Gordzevich R, Rachwalski K, Tu MM, Bordeleau E, Dooley D, Griffiths E, Zubyk HL, Brown ED, Maguire F, Beiko RG, Hsiao WWL, Brinkman FSL, Van Domselaar G, McArthur AG. 2023. CARD 2023: expanded curation, support for machine learning, and resistome prediction at the Comprehensive Antibiotic Resistance Database. Nucleic Acids Res 51:D690–d699.

