## Supplementary files for "Mutations in the riboflavin biosynthesis pathway confer resistance to furazolidone and abolish the synergistic interaction between furazolidone and vancomycin in *Escherichia coli*"

### Supplementary Figures

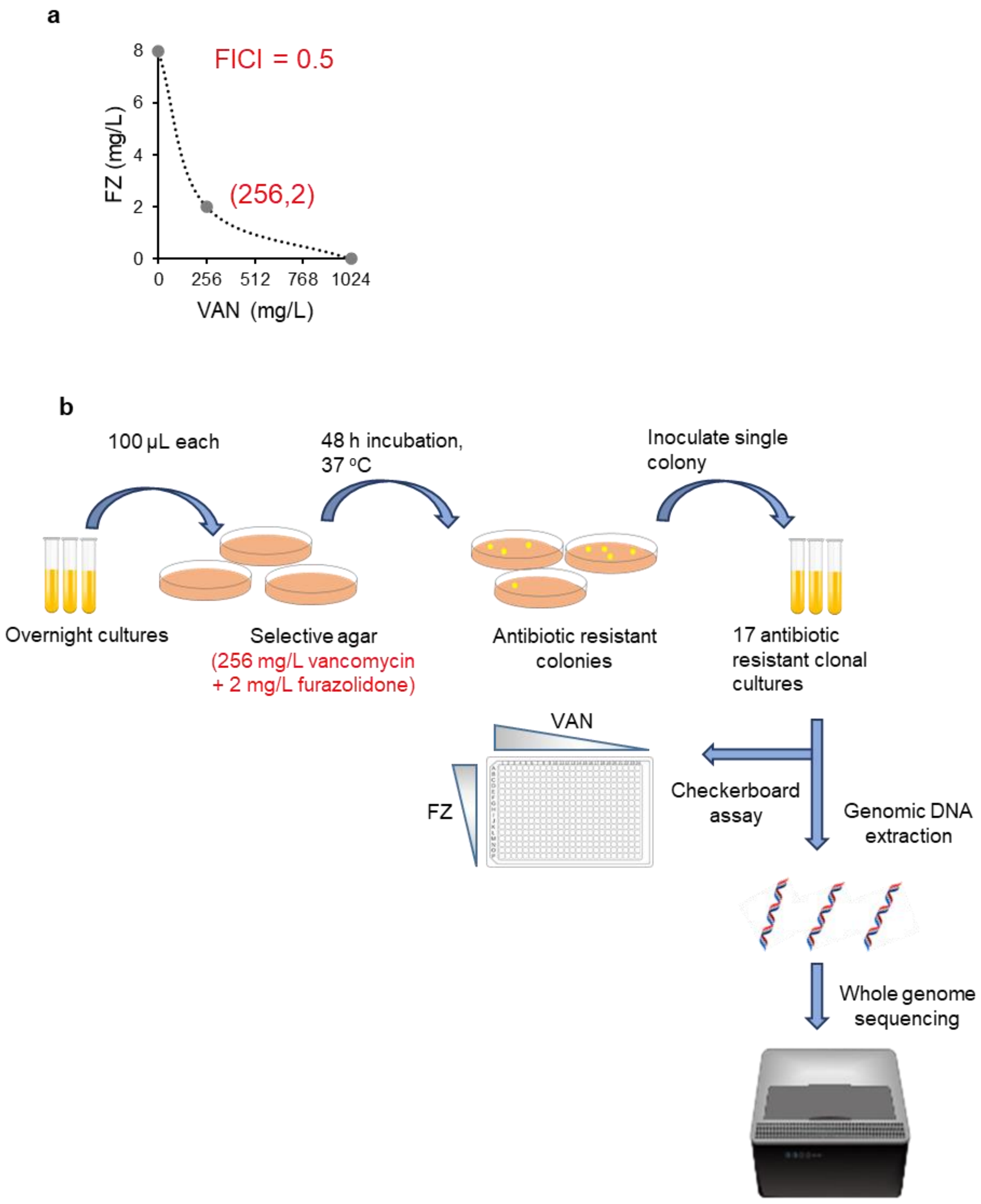


**Supplementary Figure S1.** Selecting for *E. coli* mutants resistant to the synergistic furazolidone-vancomycin combination. a) Isobologram of the agar checkerboard assay of vancomycin and furazolidone using 100 µL of overnight culture of parental strain BW25113 (PS). Each data point indicates the minimum inhibitory concentration. The synergistic interaction was maintained as shown by an FICI value of 0.5, indicating synergy. b) The workflow of isolating resistant mutants, followed by checkerboard assays to evaluate the drug interaction and genomic sequencing to identify mutations.

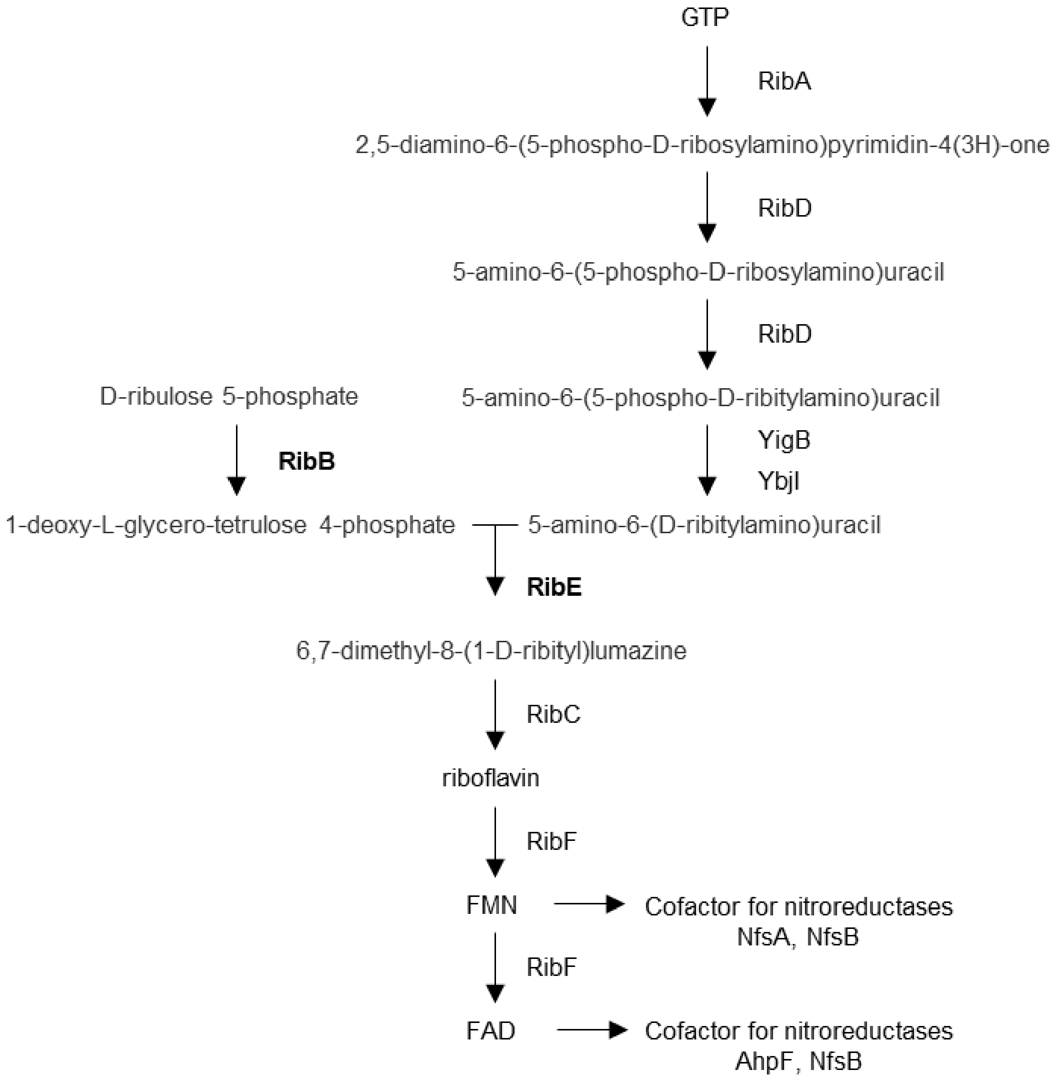


**Supplementary Figure S2:** Riboflavin biosynthesis pathway**.** Schematic illustration of the riboflavin biosynthesis pathway in *E. coli*. The name of the compounds and catalytic enzymes for each step are shown. RibB and RibE are indicated in bold (1, 2).

#### References


1. Keseler IM, Collado-Vides J, Santos-Zavaleta A, Peralta-Gil M, Gama-Castro S, Muñiz-Rascado L, Bonavides-Martinez C, Paley S, Krummenacker M, Altman T, Kaipa P, Spaulding A, Pacheco J, Latendresse M, Fulcher C, Sarker M, Shearer AG, Mackie A, Paulsen I, Gunsalus RP, Karp PD. 2011. EcoCyc: a comprehensive database of Escherichia coli biology. Nucleic Acids Res 39:D583-90.
2. Bacher A, Eberhardt S, Fischer M, Kis K, Richter G. 2000. Biosynthesis of vitamin b2 (riboflavin). Annu Rev Nutr 20:153-67.
